# Estrogen-Nitric Oxide Signaling Modulates Mitochondrial Dynamics and Endothelial Lipid Handling to Protect Against Early Atherosclerosis

**DOI:** 10.64898/2026.03.30.715353

**Authors:** Emmalie Spry, Hannah Strcula, Giovana A. Mascoli, Czarina P. Sobejana, Mirabella Zingales, Marta H. Krieger, Alessandro G. Salerno, Amarylis C. B. A. Wanschel

## Abstract

**Background:** Sex-related differences in cardiovascular disease suggest the presence of intrinsic vasoprotective mechanisms, with estrogen recognized as an important modulator of endothelial function. Building on existing evidence, the present study provides mechanistic insights into how estrogen and nitric oxide (NO) signaling regulate selective pathways of oxLDL uptake, mitochondrial dynamics, and inflammatory responses during early atherogenesis.

**Methods:** We combined an *in vitro* endothelial cell-macrophage co-culture model with *in vivo* studies in low-density lipoprotein receptor-knockout (LDLr^−/−^) mice to investigate the role of estrogen in early atherosclerotic processes. Human aortic endothelial cells (HAECs) were exposed to oxidized low-density lipoprotein (oxLDL) in the presence or absence of 17β-estradiol (E2) and the nitric oxide (NO^•^) donor S-nitroso-N-acetylcysteine (SNAC). Key outcomes included oxLDL uptake, mitochondrial oxidative stress, mitochondrial dynamics, and inflammatory signaling. *In vivo,* male and female LDLr^−/−^ mice were exposed to a short-term high-fat diet with or without SNAC treatment. Plasma lipid levels, blood pressure, aortic lesion formation, and cardiac remodeling were evaluated.

**Results:** E2 reduced oxLDL uptake and oxidative stress, effects recapitulated by SNAC; however, these responses involved distinct entry pathways, with E2 preferentially modulating lectin-like oxidized low-density lipoprotein receptor-1 (LOX-1) dependent uptake and SNAC targeting caveolae-associated mechanisms. In parallel, both E2 and SNAC reduced Scavenger Receptor Class B Type 1 (SR-BI) expression, suggesting an additional modulation on oxLDL transcytosis via this mechanism. Endothelial cells exposed to oxLDL exhibited altered mitochondrial regulatory proteins, including superoxide dismutase 2 (SOD-2), dynamin-related protein 1 (Drp-1), and optic atrophy protein 1 (OPA-1). Despite reducing oxidative stress, E2 increased the expression of adhesion molecules and enhanced monocyte adhesion in response to oxLDL exposure, particularly when combined with SNAC. Strikingly, E2 also modulated macrophage responses, increasing interleukin receptor antagonist (IL-1ra) expression and reducing GDF15, macrophage inhibitory factor (MIF), macrophage inflammatory protein 3 alfa (MIP-3α), and matrix metalloproteinase 9 (MMP-9) levels, consistent with a less pro-inflammatory macrophage profile. In vivo, HFD increased plasma lipid levels and atherosclerotic lesion area in LDLr⁻/⁻ mice, whereas SNAC partially attenuated these effects without affecting plasma lipid levels. In vivo, female LDLr−/− mice developed approximately 50% smaller aortic lesions than males, despite comparable or higher plasma lipid levels. A dyslipidemia led to increased blood pressure and a hypertensive phenotype in both males and females. SNAC treatment reduced lesion burden in both sexes and prevented diet-induced hypertension in females.

**Conclusion:** Estrogen limits early atherogenic injury by reducing endothelial uptake of oxLDL, preserving mitochondrial homeostasis, and modulating inflammatory signaling. Together, the E2 and NO pathways regulate early atherosclerosis through distinct yet complementary mechanisms, offering a potential framework for vascular-protective strategies.

## 1. INTRODUCTION

Cardiovascular disease (CVD) remains the leading cause of mortality worldwide, with well-established sex-related differences in disease onset and progression (1). Although women and men share a similar lifetime risk of coronary artery disease (CAD), disease onset in women is typically delayed, suggesting the presence of intrinsic vasoprotective mechanisms. Estrogens have been widely implicated in this protection through their effects on endothelial function, lipid metabolism, and inflammatory signaling (2). The decline in estrogen levels during menopause is associated with increased cardiovascular risk, underscoring the importance of understanding estrogen-dependent vascular mechanisms or identifying molecular pathways that may be therapeutically leveraged during the initial stages of disease.

Atherosclerosis underlies the majority of coronary artery disease (CAD) cases and is a chronic inflammatory disease of the arterial wall that develops over decades prior to clinical manifestation (3, 4). Early atherosclerosis is characterized by endothelial dysfunction in response to classical cardiovascular risk factors, including dyslipidemia, hypertension, diabetes mellitus, and smoking (5, 6), and is associated with reduced nitric oxide (NO) bioavailability and increased oxidative stress (7, 8). Disruption of endothelial redox homeostasis and mitochondrial redox imbalance promotes an oxidative environment that increases LDL transcytosis and favors its modification, leading to lesion initiation. Importantly, this initial phase represents a potentially reversible window during which targeted interventions may alter long-term cardiovascular outcomes (9–11). Although lifestyle-modifiable risk factors, such as physical inactivity, unhealthy diet, and hypertension, are broadly similar between men and women, their biological impact may differ in a sex-specific manner (12, 13). Experimental and clinical evidence indicate that estrogens exert vasoprotective effects through multiple mechanisms, including enhancement of endothelial NO synthase (eNOS) activity, suppression of oxidative stress, regulation of lipid metabolism, and modulation of inflammatory signaling (2, 14).

Despite substantial evidence supporting a protective role for estrogen signaling in atherosclerosis, key cellular mechanisms by which estrogen regulates oxLDL uptake, mitochondrial function, macrophage activation, and endothelial–immune interactions during atherogenesis remain poorly understood. While prior studies have demonstrated the vasoprotective effects of estrogen and nitric oxide, our findings provide new mechanistic insight by integrating endothelial lipid handling, mitochondrial dynamics, and immune responses, and by identifying a dissociation between endothelial activation and macrophage inflammatory signaling. Addressing these mechanisms will shed light on how estrogen-dependent pathways influence early disease and may inform future therapeutic strategies.

In the present study, we combined *in vitro* vascular cell models with *in vivo* experimental approaches to investigate how estrogen and NO signaling modulate early atherogenic processes. Specifically, we examined their effects on oxLDL uptake pathways, mitochondrial dynamics, endothelial activation, and macrophage inflammatory responses to identify pathway-specific mechanisms contributing to early atherosclerotic development.

## 2. MATERIALS AND METHODS

### 2.1 Human Aortic Endothelial Cell Culture (HAECs)

HAECs, the culture endothelial basal medium (EBM, CC-3121), and endothelial cell growth supplements (EGM, CC-4133) were purchased from Lonza (Walkersville, MD, USA). The cells were grown in EBM containing 15% fetal bovine serum (FBS, R&D, Flowery Branch, GA, USA) at 37◦C in a humidified 5% CO2 incubator and used for experiments between passages 3 and 11. HAECs were allowed to grow until 100% confluence with or without 17-βestradiol (E2) in human serum BCR-1348 (Community Bureau of Reference, European Commission, Brussels, Belgium) and treated with human oxidized low-density lipoprotein (Invitrogen L34357) and S-nitroso-N-acetylcysteine (SNAC) synthesized as described by Ricardo et al. (15) using reagent-grade chemicals from Alfa-Aesar (Nitrite #14244) and Millipore (N-Acetylcysteine-L-cysteine #106425). Forty-eight hours after treatments, HAECs were harvested for subsequent analyses.

### 2.2 Human Monocytic Cells

Human THP-1 monocytes were purchased from ATCC (Manassas, VA, USA). These cells were cultured in RPMI-1640 medium supplemented with 10% (v/v) heat-inactivated FBS, 100 U/mL penicillin, 100 µg/mL streptomycin, and 2 mM L-glutamine (all Invitrogen). The culture was maintained at 37 ◦C in a humidified atmosphere containing 5% CO2.

### 2.3 Preparation of Fluorescent THP-1 Cells

To visualize THP-1 cells adhering to the HAECs, fluorescent THP-1 cells were prepared as described by the manufacturer (C2925, Invitrogen). In brief, THP-1 cells were labeled with a fluorescent dye by incubating 1.0 × 106 cells in 10 mL of FBS-free RPMI-1640 and 10 µl of a cell-tracker (final concentration 5 µM) at 37 ◦C for 45 min. Dye loading was stopped by centrifugation, and cells were resuspended in fresh EBM medium.

### 2.4 Adhesion Experiment of THP-1 Cells/Monocyte-EC adhesion

To investigate the effects of E2 and SNAC on the adhesion of THP-1 cells to the endothelial monolayer, HAECs were plated on Lab-Tek slides (Thermo Scientific, Rochester, NY, USA), and the monolayers were treated with oxidized low-density lipoprotein. Human THP1 monocytes previously incubated with 5 µM Cell-Tracker Green (CMFDA) (Invitrogen, Eugene, OR, USA) and resuspended in 500 µl of fresh EBM medium were placed into each well of a Lab-Tek slide and incubated at 37 °C for 30 min. Next, the THP-1 cell suspension in each well was discarded, and the HAEC monolayer was gently washed three times with 500 µL of phosphate-buffered saline (PBS, pH 7.3) in order to remove non-adherent THP-1 cells. Adherent fluorescence-labeled THP-1 cells were observed under an EVOS FL Microscope (Life Technologies, Carlsbad, SD, USA) and quantified.

### 2.5 Immunofluorescence Microscopy

For LDLR internalization imaging in HAECs, the cells were plated on chamber slides (Lab-Tek II, Thermo Scientific) and allowed to grow to 100% confluence with or without 17β-estradiol (E2) in human serum, then incubated with diI-LDL (Invitrogen, L34358) for 30 min and washed with cold PBS. The cells were fixed with 4% PFA in PBS. Cells treated with diI-LDL were incubated with DAPI (D-9542, Sigma Aldrich) for 10 min. After washing, cells were mounted in a fluorescence mounting medium (H-1000, Vector). Images were collected with an EVOS FL Microscope (Life Technologies, Carlsbad, SD, USA) and quantified. The manufacturer’s software was used for data acquisition, and ImageJ for fluorescence profiles.

### 2.6 Measurement of Mitochondrial Superoxide Production in HAECs

HAECs were allowed to grow until 100% confluence with or without 17βestradiol (E2) in human serum and treated with oxidized low-density lipoprotein and or S-nitroso-N-acetylcysteine on 8-well LabTek slides. Cells were pre-incubated with 5 µM of the mitochondrial superoxide indicator MitoSOX (Molecular Probes, Invitrogen - M36008) at 37 ◦C for 30 min. Cells were mounted onto glass slides with Mounting Medium Vectashield (Vector Laboratories). Images of cells were acquired using an EVOS FL Microscope (Life Technologies, Carlsbad, SD, USA) with 10× magnification. Each well was imaged using transmitted light with GFP- and RFP-specific LED cubes, followed by measurement of the average cell area of at least 10 cells per field using ImageJ software.

### 2.7 Western Blot

Cells were homogenized in RIPA buffer (Cell Signaling c#9803S) supplemented with protease inhibitor and phosphatase inhibitor cocktail (Abcam ab5872S). Protein lysates were boiled with Laemmli Sample Buffer (Bio-Rad, Hercules, CA) containing 355 mM b-mercaptoethanol, and aliquots with equal amounts of protein were loaded on 7.5% or 4%–20% SDS-PAGE gels (Bio-Rad, Hercules, CA) and run. Subsequently, separated proteins were transferred onto PVDF membranes (Bio-Rad, Hercules, CA). PVDF membrane blots were blocked in 5% BSA in TBS-T (20 mM Tris, 0.16 M NaCl, and 0.10% Tween-20, pH 7.4) for 1 h at RT, and incubated overnight at 4°C with rabbit anti-SOD2 antibodies (1:1,000, ab13533, Abcam); rabbit anti-DRP-1 Ser616 antibodies (1:1,000, #3455S, Cell Signaling Technology, Danvers, MA); rabbit anti-OPA-1 antibodies (1:1,000, #2382, Cell Signaling Technology, Danvers, MA); mouse anti-SRA (MSR1) antibodies (1:1,000, #17275S, Cell Signaling Technology, Danvers, MA); rabbit anti-LOX-1 antibodies (1:1,000, ab14427, Abcam); rabbit anti-Caveolin-1 antibody (1:1,000, ab2910, Abcam); rabbit anti-LDL receptor antibody (1:1,000, ab30532, Abcam); rabbit anti-SR-BI antibodies (1:1,000, ab137829, Abcam), and rabbit anti-GAPDH antibodies (1:1,000, #G9545, Sigma-Aldrich); rabbit anti-HSP-90 antibodies (1:1,000, #4874S, Cell Signaling Technology, Danvers, MA) in 5% BSA in TBS-T. Anti-rabbit IgG (#7074S, Cell Signaling Technology, Danvers, MA) and anti-mouse IgG (#7076S, Cell Signaling Technology, Danvers, MA) were used as secondary antibodies. The bands were detected using the Odyssey CLx Imaging System (LI-COR).

### 2.8 THP-1 macrophages Induction

To differentiate THP-1 cells into THP-1 macrophages (THP-1m) for our co-culture experiment, we seeded cells at densities of 2 × 10 ^5 and 1 × 10^6 cells/well in six-well plates. PMA (Sigma-Aldrich, Saint Louis, MO, USA; cat#: 16561298) was added at doses of 100 ng/mL. After 48 h of PMA stimulation, THP-1m was obtained.

### 2.9 THP-1m Culture

Following differentiation, THP-1-derived macrophages (THP-1m) were washed to remove PMA and maintained in PMA-free medium for 24 h. Cells were subsequently co-cultured with HAECs using Transwell inserts and exposed to oxLDL, E2, or SNAC for 24 h. After incubation, cells were harvested and homogenized in RIPA buffer as described above, and cytokine levels were analyzed using the Proteome Profiler Human XL Cytokine Array Kit (R&D Systems, Cat. No. ARY022B).

### 2.10 Animals and Diet

Experiments used 3-month-old (24 ± 3 g, n = 120) male LDLr^−/−^mice from Jackson Laboratory (Bar Harbor, ME) and wild-type mice (strain C57BL/6/uni) obtained from the animal facilities at UNICAMP. The LDL-Rtm1Her mutant strain was developed, and the129-derived AB1 ES cell line was used. The strain was backcrossed to C57BL/6J mice for 10 generations. The transgenic mice used here, LDLr^−/−^and the C57BL/6/Uni strain maintained in our unit were analyzed to guarantee its use as a control group. For this purpose, DNA samples from LDLr^−/−^ (stock 002207), C57BL/6/Uni, and 129/SvPas (Steel substrain derived) mice were analyzed using a panel of microsatellite markers. The results showed no differences between the C57BL/6/Uni and LDLr^−/−^ congenic strains used in this study, and identified polymorphic regions in the 129/SvPas strain. This suggests that the LDLr^−/−^ strain is congenic with the C57BL/6 background. The experimental protocols were approved by the Institutional Committee for Ethics in Animal Experimentation (IACUC: 2025-1112 and CEEA/IB - UNICAMP, protocol no. 521-1) in agreement with the guidelines of the Brazilian College for Animal Experimentation (COBEA). The mice were randomly allocated to one of the 4 groups described below and received ad libitum access to diet and water. (1) C57BL/6/Uni used as wild-type group (WT) [daily dose, 0.1 mL of phosphate buffered saline (PBS) IP], which received standard diet (Nuvital CR1); (2) LDLr^−/−^ control group (S) (daily dose, 0.1 mL PBS IP), which received a standard diet (Nuvital CR1); (3) Hypercholesterolemic LDLr^−/−^ group (Chol) (daily dose, 0.1 mL PBS IP), which received an atherogenic diet (high-cholesterol diet: 20% fat, 1.25% cholesterol, 0.5% cholic acid); and (4) similarly handled hypercholesterolemic LDLr^−/−^ mice were given SNAC (Chol + SNAC group) (daily dose, 0.51 μmol/kg IP). (15, 16). After 15 days, mice were anesthetized with a mixture of xylazine and ketamine (6 mg/kg and 40 mg/kg, respectively, IP). The measurements were performed in a blinded fashion.

### 2.11 Resting Blood Pressure and Heart Rate Measurements

Tail-cuff blood pressure and heart rate were measured in conscious mice before treatment with SNAC between 10 am and 12 am, using a computerized tail-cuff Kent Scientific (XBP 1000) system. The first 6 days of measurements were primarily for training. Data collected during these days were not used for calculations but to assess whether reliable flow waveforms could be obtained in each mouse. On the day of the recordings, sets of 30 measurements were recorded. On average, 20 to 30 blood pressure value measurements were computed for each mouse (17).

### 2.12 Plasma lipoproteins

Plasma from C, HC or SNAC + HC mice was obtained by centrifugation (12,000 rpm, 15 min) of blood collected in heparinized tubes via the retro orbital plexus. Total cholesterol and triglyceride were determined by enzymatic colorimetric kits (Wako Chemicals). Individual plasma samples (200 μl) were fractionated by FPLC using an HR10/30 Superose6 column (Amershan/Pharmacia Biotech), balanced with Tris–saline buffer, pH 7.4. Total cholesterol was determined enzymatically in each FPLC fraction. Results (mg/dL) are from 3 mice/group.

### 2.13 Histological analysis of ascending proximal aorta atherosclerotic lesions

Mice were anesthetized and their hearts were perfused in situ with phosphate-buffered saline (PBS) followed by 10% PBS-buffered formaldehyde, after which they were excised and fixed in 10% formaldehyde for at least 2 days. The hearts were then embedded sequentially in 5, 10, and 25% gelatin. Processing and staining were carried out according to (18). The lipid-stained lesions were quantified as described by (19) using Image Pro Plus software (version 3.0) for image analysis (Media Cybernetics, Silver Spring, MD). The slides were read by an investigator who was unaware of the treatments. The area of the lesions was expressed as the sum of the lesions across seven 10 μM sections, 80 μM apart, for a total aorta length of 480 μM. Because several other studies revealed a predilection for the development of lesions in the aortic root, the segment that was chosen for analysis extended from beyond the aortic sinus up to the point where the aorta first becomes rounded. Results from six mice/group are expressed in μM^2^.

### 2.14 Statistical Analyses

Mean values between the two experimental groups were compared by an unpaired two-tailed Student t-test (SigmaPlot 15). Unless stated otherwise, data are expressed as mean ± standard error of the mean (SEM). p < 0.05 was set as the statistical significance level.

## 3. RESULTS

### 3.1 In Vitro Studies: Estrogen prevents oxLDL uptake and transcytosis in endothelial cells

To investigate whether estrogen protects endothelial cells from early atherogenic injury, HAECs were exposed to Dil-labeled oxidized low-density lipoprotein (Dil-oxLDL) in the presence or absence of 17β-estradiol (E2) and/or the NO donor S-nitroso-N-acetylcysteine (SNAC). Representative fluorescence images showed a robust intracellular Dil-oxLDL signal after oxLDL exposure, whereas E2 treatment reduced oxLDL accumulation in endothelial cells. SNAC treatment produced a similar reduction, and co-treatment with E2 + SNAC did not further enhance the protective effect (**Figure 1A**). Quantitative analysis confirmed that E2 significantly attenuated oxLDL uptake, supporting the hypothesis that estrogen prevents early-stage atherosclerosis. SNAC similarly reduced Dil-oxLDL uptake; however, combined treatment with E2 and SNAC did not further reduce oxLDL uptake beyond that of either intervention alone (**Figure 1B**). However, E2 and SNAC differentially regulate oxLDL uptake pathways in endothelial cells. The mechanism seems to involve distinct entry pathways, with E2 primarily downregulating LOX-1 and enhancing LDL receptor expression, suggesting a shift toward physiological lipid uptake. SNAC predominantly targets caveolae-associated mechanisms through marked reduction of caveolin-1 expression. In parallel, both E2 and SNAC reduced Scavenger Receptor Class B Type 1 (SR-BI) expression, suggesting an additional modulation on oxLDL transcytosis via this mechanism (**Figure 1C**).

**Figure 1.**
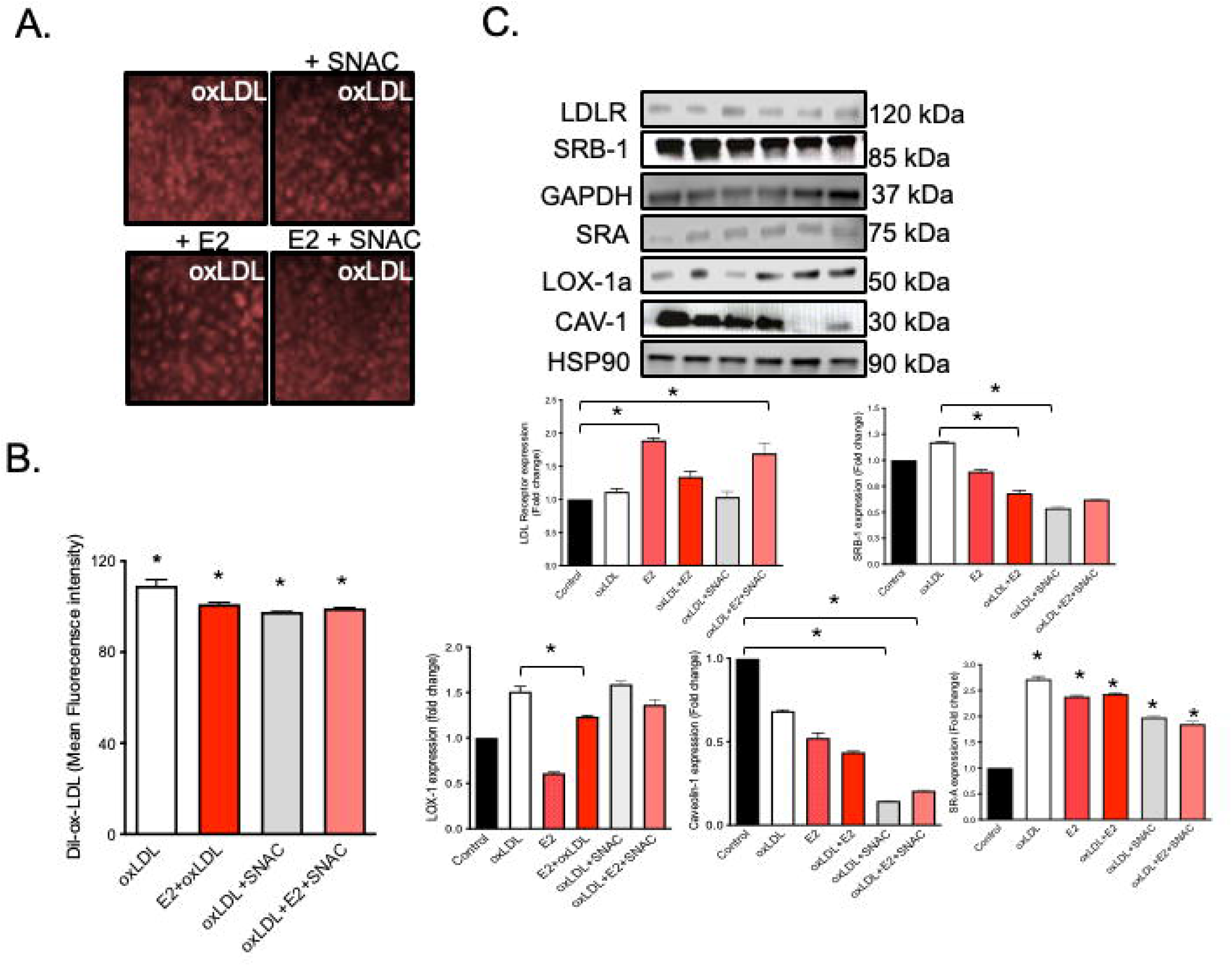
Estrogen and nitric oxide signaling differentially regulate oxLDL uptake pathways in endothelial cells. (A) Representative fluorescence images of DiI-oxLDL uptake in human aortic endothelial cells (HAECs) under oxLDL exposure with or without 17β-estradiol (E2), S-nitroso-N-acetylcysteine (SNAC), or combined treatment. (B) Quantification of DiI-oxLDL fluorescence intensity, showing reduced oxLDL uptake following E2 and SNAC treatment. (C) Representative Western blots and corresponding densitometric analyses of proteins involved in lipid uptake pathways, including LDL receptor (LDLR), scavenger receptor class B type 1 (SR-BI), scavenger receptor A (SR-A), lectin-like oxidized LDL receptor-1 (LOX-1), and caveolin-1 (Cav-1). E2 increased LDLR expression and reduced LOX-1 levels, while SNAC markedly reduced Cav-1 expression. Both treatments decreased SR-BI expression, whereas SR-A showed minimal changes. Data are expressed as mean ± SEM. *p < 0.05 versus Control or oxLDL.

### 3.2 Estrogen suppresses oxLDL-induced mitochondrial oxidative stress

OxLDL exposure significantly increased mitochondrial superoxide (O_2_^−•^) production in endothelial cells, as measured by MitoSOX Red fluorescence. Estrogen treatment significantly reduced mitochondrial superoxide generation under oxLDL conditions (**Figure 2A**). Western blot analysis further suggested that E2 enhances the expression of the mitochondrial antioxidant enzyme superoxide dismutase 2 (SOD2), supporting a role for estrogen in preserving mitochondrial redox homeostasis during early atherogenic stress. Notably, combined E2 and SNAC treatment further increased SOD2 expression, suggesting enhanced protection against oxidative stress. Together, these findings indicate that estrogen and nitric oxide donor treatment strengthen mitochondrial antioxidant defenses during oxLDL-induced stress (**Figure 2B**).

**Figure 2.**
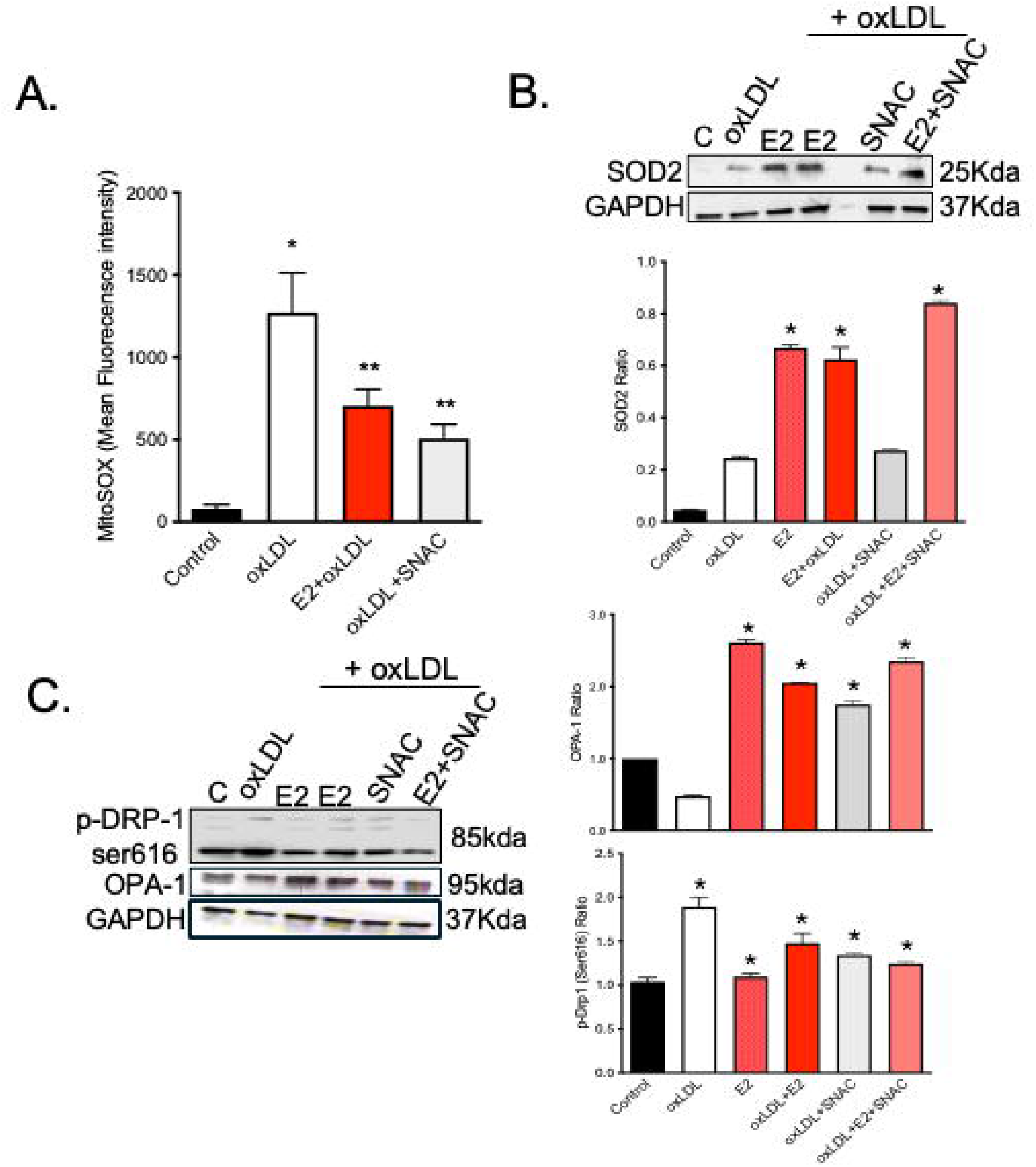
Estrogen and nitric oxide signaling modulate mitochondrial oxidative stress and dynamics in endothelial cells. (A) Quantification of mitochondrial superoxide production in human aortic endothelial cells (HAECs) measured by MitoSOX fluorescence under control conditions and following oxLDL exposure with or without 17β-estradiol (E2) or S-nitroso-N-acetylcysteine (SNAC). (B) Representative Western blots and densitometric analysis of superoxide dismutase 2 (SOD2) expression under the indicated conditions. (C) Representative Western blots and quantification of mitochondrial dynamics-related proteins, including phosphorylated dynamin-related protein 1 at Ser616 (p-Drp1 Ser616) and optic atrophy protein 1 (OPA1), in HAECs exposed to oxLDL with or without E2, SNAC, or combined treatment. Data are expressed as mean ± SEM. *p < 0.05 versus Control or ** versus oxLDL.

### 3.3 Estrogen selectively attenuates mitochondrial fission signaling

To determine whether estrogen influences mitochondrial quality control pathways, proteins involved in mitochondrial fission and fusion were examined. OxLDL increased Drp1 phosphorylation at Ser616, a modification associated with mitochondrial fission and fragmentation. E2 reduced the oxLDL-induced increase in phospho-Drp1 (Ser616), indicating suppression of stress-associated mitochondrial fission signaling. SNAC treatment also attenuated Drp1 Ser616 phosphorylation, and combined E2 + SNAC treatment did not produce a clear additional reduction, suggesting an overlapping protective mechanism (**Figure 2C**).

In contrast, OPA1 expression, a marker of mitochondrial fusion and cristae integrity, was modestly increased by oxLDL alone, suggesting an early compensatory protective response. This effect was further enhanced by E2, SNAC, and combined E2 + SNAC treatment, indicating promotion of fusion-associated mitochondrial remodeling under these conditions (**Figure 2C**). Collectively, these findings suggest that estrogen not only limits pro-fission signaling but also promotes mitochondrial fusion remodeling.

### 3.4 Estrogen modulates endothelial inflammatory signaling and monocyte adhesion

Endothelial cells exposed to oxLDL were co-cultured with THP-1–derived macrophages to evaluate endothelial–immune interactions during the early stages of atherogenesis. Surprisingly, estrogen treatment increased monocyte adhesion to the endothelium under oxLDL conditions (**Figure 3A-B**). THP-1-derived macrophages (THP-1m) exposed to oxLDL increased the expression of adhesion molecules and chemokines associated with leukocyte recruitment, including ICAM-1, MCP-1, and IL-8. Furthermore, THP-1m cells exposed to estrogen exhibited enhanced expression of the anti-inflammatory mediator IL-1 receptor antagonist (IL-1Ra) and a reduction in the pro-inflammatory cytokines growth differentiation factor-15 (GDF-15), MIF, MIP-3α, and MMP-9, suggesting coordinated modulation of inflammatory signaling during early lesion formation (**Figure 3C-J**).

**Figure 3.**
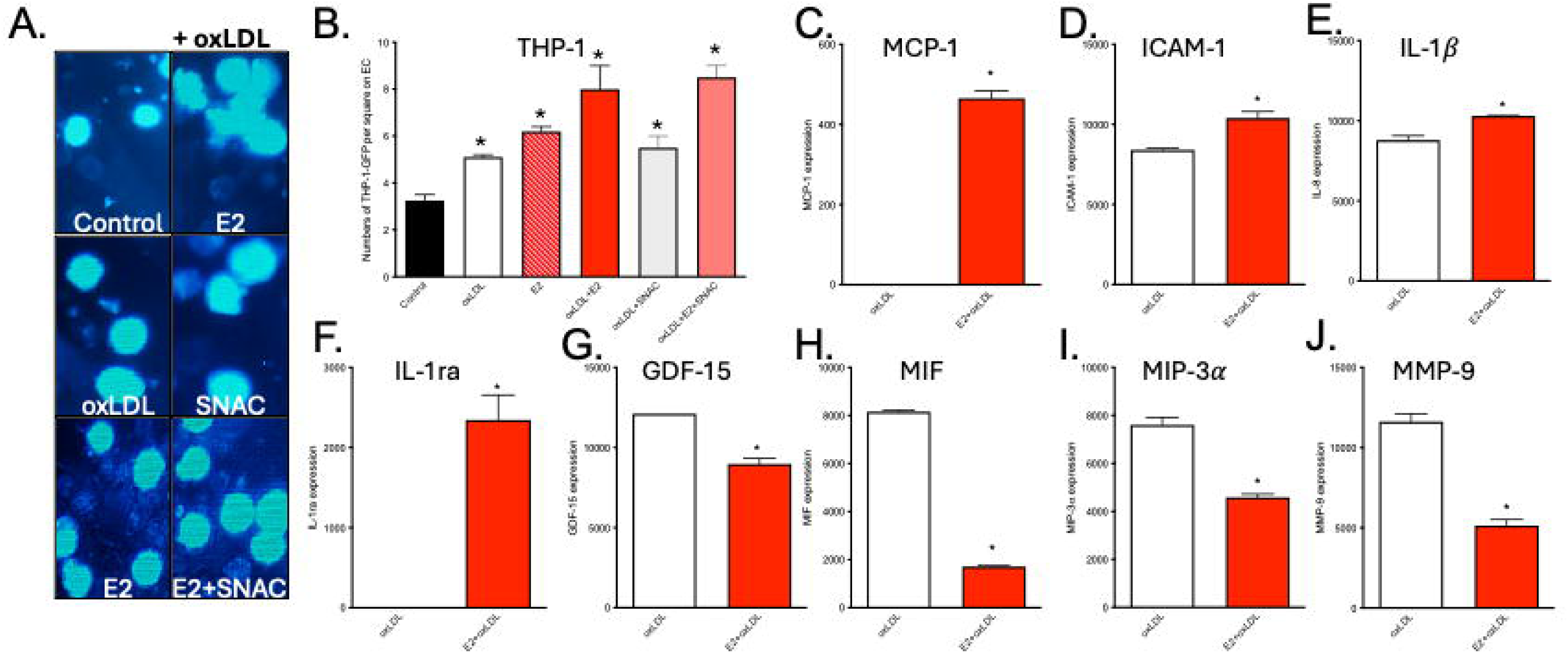
Estrogen modulates endothelial–monocyte interactions and macrophage inflammatory signaling. (A) Representative fluorescence images of THP-1 monocyte adhesion to human aortic endothelial cells (HAECs) under control conditions and following oxLDL exposure with or without 17β-estradiol (E2), S-nitroso-N-acetylcysteine (SNAC), or combined treatment. (B) Quantification of adherent THP-1 cells per endothelial cell monolayer under the indicated conditions. (C–E) Expression levels of endothelial inflammatory markers, including MCP-1, ICAM-1, and IL-1β, in response to oxLDL with or without E2 treatment. (F–J) Expression of macrophage-derived inflammatory mediators, including IL-1Ra, GDF-15, macrophage migration inhibitory factor (MIF), macrophage inflammatory protein-3α (MIP-3α/CCL20), and matrix metalloproteinase-9 (MMP-9), measured in THP-1–derived macrophages following co-culture under the indicated conditions. Data are expressed as mean ± SEM. *p < 0.05 versus oxLDL.

Collectively, these in vitro findings indicate that estrogen protects against key early atherogenic processes, including endothelial lipid uptake, mitochondrial oxidative stress, and dysregulated mitochondrial dynamics, while dynamically modulating inflammatory signaling and immune cell interactions at the vascular interface.

### 3.5 Female LDLr−/− mice exhibit reduced early lesion formation compared with males

To determine whether the vascular protection observed in vitro was reflected in vivo, early atherogenesis was evaluated in male and female LDL receptor-deficient (LDLr−/−) mice exposed to a short-term hypercholesterolemic diet. After 15 days, lipid-rich lesions were detected in the proximal aorta in both sexes; however, female mice developed substantially smaller lesions than male mice under the same dietary conditions. Lesion area in females was reduced by approximately 50%, indicating pronounced sexual dimorphism during the earliest stage of atherosclerotic lesion development (**Figure 4G**).

**Figure 4.**
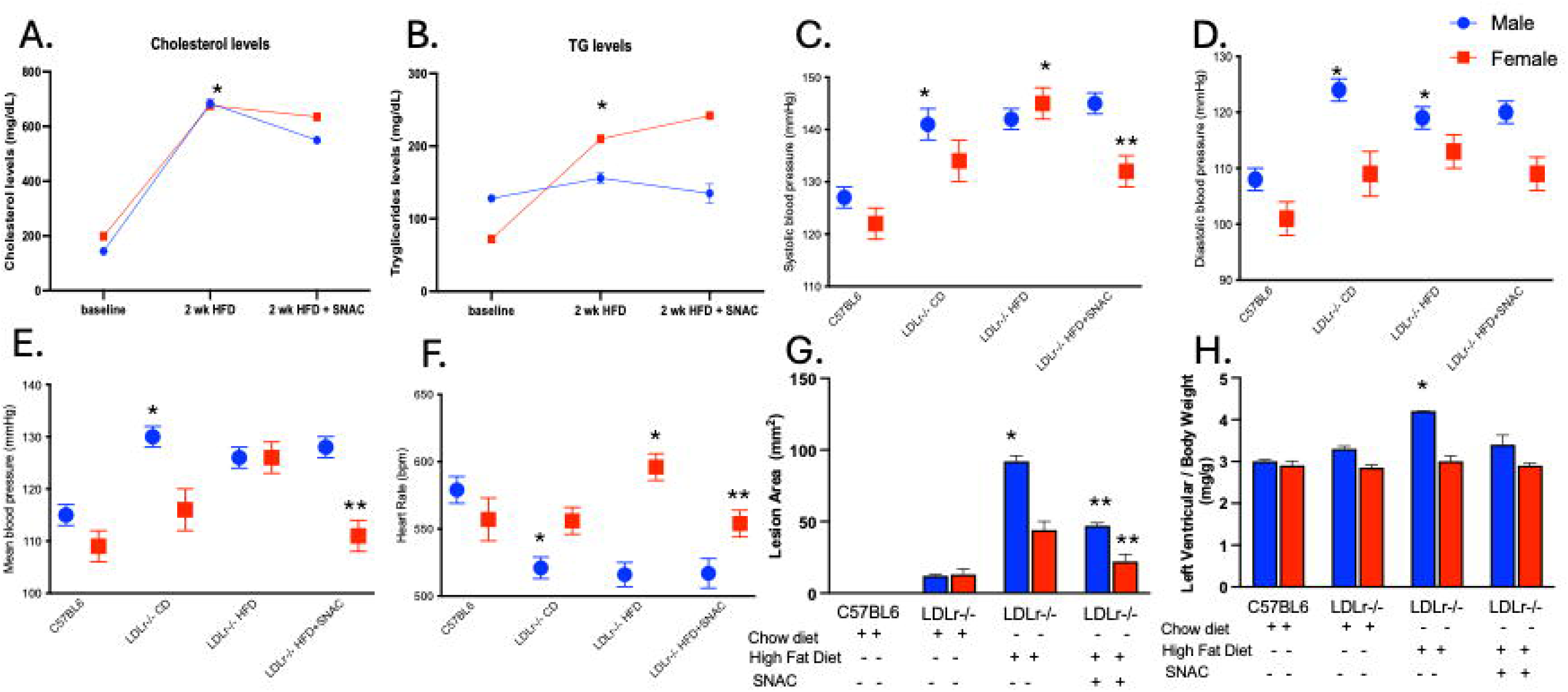
Sex-specific differences and nitric oxide–dependent effects on early atherosclerosis and cardiovascular parameters in LDLr⁻/⁻ mice. (A) Plasma cholesterol levels measured at baseline, after 2 weeks of high-fat diet (HFD), and after HFD with S-nitroso-N-acetylcysteine (SNAC) treatment in male and female mice. (B) Plasma triglyceride (TG) levels under the same experimental conditions. (C–E) Systolic (C), diastolic (D), and mean arterial blood pressure (E) measured in control (C57BL6), LDLr⁻/⁻ mice on chow diet, LDLr⁻/⁻ mice on HFD, and LDLr⁻/⁻ mice on HFD treated with SNAC, separated by sex. (F) Heart rate measurements across experimental groups. (G) Quantification of atherosclerotic lesion area in the proximal aorta under the indicated dietary and treatment conditions. (H) Left ventricular weight normalized to body weight across experimental groups. Data are expressed as mean ± SEM. *p < 0.05 versus CT and ** versus LDLr⁻/⁻ HFD.

This protective phenotype occurred despite hypercholesterolemic challenge, suggesting that endogenous female sex hormones may limit lesion initiation through direct vascular mechanisms rather than solely through changes in circulating lipids.

### 3.6 NO donor therapy attenuates early lesion development in both sexes

SNAC reduces early atherosclerotic lesion burden in both sexes.

Given the in vitro evidence that nitric oxide-related signaling contributes to endothelial protection, the effect of SNAC treatment was assessed in vivo. SNAC reduced aortic lesion area in both male and female LDLr−/− mice fed the hypercholesterolemic diet. The magnitude of lesion reduction was similar in both sexes, approximately 50% relative to untreated hypercholesterolemic animals, indicating that augmentation of nitric oxide bioavailability limits early lesion formation independently of sex (**Figure 4G**).

Notably, SNAC treatment diminished the sex difference in lesion burden by reducing lesion size in males to levels closer to those observed in females. These data support nitric oxide signaling as a key atheroprotective pathway during lesion initiation.

### 3.7 Atheroprotection in females is uncoupled from plasma lipid levels

To determine whether reduced lesion formation in females was secondary to a more favorable systemic metabolic profile, plasma lipid levels were compared between groups. Despite developing smaller lesions, female LDLr−/− mice displayed elevated cholesterol and triglyceride levels under hypercholesterolemic conditions (**Figure 4A-B**). Moreover, SNAC treatment reduced lesion burden without normalizing plasma lipid concentrations. Together, these findings indicate that the observed vascular protection in females, as well as the protective effect of SNAC, is not primarily attributable to lipid lowering. These results instead support the concept that estrogen- and nitric oxide-dependent mechanisms act directly at the vascular wall to limit endothelial dysfunction and early lesion formation.

### 3.8 Sex-specific regulation of blood pressure during early atherogenesis

Hypercholesterolemic diet induced a significant increase in systolic, diastolic, and mean arterial pressure in female LDLr−/− mice, consistent with early endothelial dysfunction (**Figure 4C-E**). Notably, SNAC treatment completely prevented diet-induced hypertension in females, restoring blood pressure values to those observed in wild-type and control LDLr−/− mice. In contrast, male LDLr−/− mice exhibited a hypertensive phenotype even under control diet conditions, and SNAC treatment failed to prevent elevated blood pressure, suggesting distinct mechanisms of blood pressure regulation between sexes, potentially involving androgen-dependent pathways.

### 3.9 Females are protected from adverse cardiac remodeling during early disease

Because sustained hemodynamic stress can promote structural remodeling of the heart, parameters of cardiac remodeling were evaluated. Male LDLr−/− mice developed left ventricular hypertrophy accompanied by reduced heart rate, consistent with early adverse cardiac remodeling (**Figure 4F-H**). In contrast, female mice did not exhibit comparable hypertrophic changes or bradycardia during the same experimental period. These findings indicate that female mice are relatively protected from early cardiac remodeling during diet-induced vascular stress. Together with the observed differences in blood pressure regulation and lesion formation, these results suggest that endogenous estrogen signaling may confer protection against early cardiovascular dysfunction during the initial stages of atherogenesis.

## 4. DISCUSSION

The present study demonstrates that estrogen signaling protects against early atherogenic injury by coordinating the regulation of endothelial lipid handling and mitochondrial oxidative stress, and by modulating inflammatory signaling. Consistent with cellular findings, female LDLr^−/-^ mice developed significantly smaller early atherosclerotic lesions compared with males (20). Importantly, pharmacological enhancement of nitric oxide bioavailability with the NO donor S-nitroso-N-acetylcysteine reproduced several of these protective effects, but via distinct endothelial entry pathways, supporting the concept that estrogen-NO signaling represents vascular protection during the initiation of atherosclerosis (21, 22).

Estrogen significantly attenuated oxLDL internalization in human aortic endothelial cells. Our findings further indicate that estrogen and nitric oxide signaling regulate oxLDL uptake through distinct but complementary endothelial pathways involving LOX-1 and caveolae-mediated transport. LOX-1 is a major receptor responsible for oxLDL internalization and is strongly associated with endothelial dysfunction, oxidative stress, and pro-inflammatory signaling during early atherogenesis. In this study, E2 reduced LOX-1 expression, consistent with prior reports demonstrating that estrogen suppresses LOX-1–dependent oxLDL uptake and downstream inflammatory activation. This reduction likely contributes to decreased oxLDL internalization and attenuation of endothelial oxidative stress.

In contrast, SNAC markedly reduced caveolin-1 expression, suggesting modulation of caveolae-dependent lipid transcytosis pathways. Caveolae are key structures involved in endothelial LDL and oxLDL trafficking across the vascular wall, and their disruption has been associated with reduced lipoprotein transcytosis and atheroprotection. These findings suggest that while estrogen primarily limits oxLDL uptake through receptor-mediated mechanisms involving LOX-1, nitric oxide signaling preferentially targets caveolae-associated transport pathways. Together, these results support a model in which estrogen and NO signaling coordinately reduce pathological lipid entry into the vascular wall by targeting distinct but convergent mechanisms, thereby limiting early endothelial dysfunction and atherogenic progression.

Estrogen has been shown to inhibit LDL transcytosis across the endothelium through mechanisms involving G-protein-coupled estrogen receptor (GPER) signaling and modulation of SR-BI–dependent pathways (23). Consistent with this, our data demonstrate that E2 increases LDLR expression while reducing LOX-1 and SR-BI levels in endothelial cells, suggesting a shift away from scavenger receptor–associated oxLDL handling and pro-atherogenic lipid trafficking toward a more regulated lipoprotein uptake profile. Given that LOX-1 is a key mediator of oxLDL internalization and pro-atherogenic signaling, its downregulation by E2 likely contributes to reduced lipid-induced oxidative and inflammatory stress. In parallel, decreased SR-BI expression may reflect reduced transcytosis of modified lipoproteins across the endothelium. Together, these findings support a model in which estrogen preserves endothelial homeostasis by limiting pathological lipid entry into the vascular wall while favoring more regulated lipoprotein handling during early dyslipidemia (24, 25).

In the present study, estrogen treatment reduced mitochondrial O ^−•^ levels and increased expression of the mitochondrial antioxidant enzyme SOD2. This ER-α dependent upregulation of MnSOD has been previously documented in endothelial cells, where it detoxifies superoxide and prevents mitochondrial damage (26–28). SNAC treatment further enhanced SOD2 expression, suggesting that N-acetylcysteine contributes to mitochondrial antioxidant defense (29). By preserving mitochondrial redox homeostasis, estrogen and SNAC may therefore limit the amplification loops of oxidative stress that drive endothelial injury (30, 31).

Estrogen influenced mitochondrial structural dynamics by attenuating phosphorylation of the mitochondrial fission protein Drp1 at Ser616 while increasing the expression of the fusion protein OPA1. Maintenance of mitochondrial network integrity is a critical determinant of endothelial resilience (32). Consistent with our findings, estrogen signaling has been shown to restore mitochondrial fission–fusion balance in vascular cells, thereby preventing pathological mitochondrial fragmentation (33).

Our co-culture findings suggest that estrogen not only modulates vascular endothelial function but also reshapes monocyte and macrophage behavior. Estrogen-and SNAC-treated THP-1 monocytes exhibited enhanced migration and adhesion to the endothelial monolayer under oxLDL conditions. Despite promoting early monocyte recruitment, estrogen drove macrophages toward a less inflammatory phenotype characterized by increased expression of the anti-inflammatory mediator IL-1Ra and reduced levels of GDF-15, MIF, MIP-3α/CCL20, and MMP-9.

Although GDF-15 is commonly induced under cellular stress and has been associated with protective and anti-inflammatory functions (34, 35), its reduction here may reflect broader suppression of stress-responsive inflammatory signaling by estrogen. Notably, estrogen exposure increased IL-1Ra while reducing mediators associated with leukocyte recruitment, inflammatory activation, and plaque destabilization. IL-1Ra functions as an endogenous inhibitor of IL-1 signaling and has been implicated in limiting lesion progression and macrophage-driven vascular remodeling. In contrast, MIF promotes monocyte recruitment and inflammatory activation, CCL20/CCR6 signaling contributes to leukocyte trafficking to the vascular wall, and MMP-9 is linked to extracellular matrix degradation and plaque vulnerability.

Together, these findings suggest that estrogen may permit early leukocyte recruitment while limiting downstream macrophage-mediated inflammatory and proteolytic amplification. These observations support a context-dependent model in which estrogen preserves vascular immune surveillance during early atherogenesis while restraining progression toward a more unstable lesion phenotype.

Importantly, female LDLr-/- mice developed smaller aortic lesions compared with males after a short-term hypercholesterolemic diet (36). Notably, this protection occurred despite elevated plasma lipid levels in females, indicating that reduced lesion formation was not primarily driven by differences in systemic lipid metabolism (37, 38). Instead, these observations support the concept that endogenous estrogen confers direct vascular protection in females (26, 39), whereas the administration of the NO donor SNAC confers significant additional protection against lesion size reduction in both sexes (6, 21). In addition to reducing lesion burden, sex-specific differences were also observed in hemodynamic and cardiac parameters. Male and female LDLr⁻/⁻ mice exhibited elevated systolic, diastolic, and mean arterial pressure, consistent with early endothelial dysfunction and increased vascular resistance under hypercholesterolemic conditions. SNAC treatment attenuated blood pressure in females, consistent with nitric oxide–mediated vasodilation and improved endothelial function.

Alterations in heart rate further reflected these differences, with males displaying reduced heart rate under hypercholesterolemic conditions, suggestive of early autonomic or cardiovascular dysfunction, whereas females exhibited a preserved cardiac profile. Consistent with these hemodynamic findings, male LDLr⁻/⁻ mice developed increased left ventricular mass, indicative of early cardiac remodeling in response to pressure overload. In contrast, female mice were protected from left ventricular hypertrophy, further supporting a cardioprotective role of estrogen signaling. Together, these findings indicate that estrogen and nitric oxide signaling not only limit early atherogenic lesion formation but also preserve vascular function and prevent early hemodynamic and cardiac maladaptation during the initial stages of disease.

## Conclusion

These findings demonstrate that estrogen limits early atherogenic injury by reducing endothelial uptake of oxLDL, preserving mitochondrial homeostasis, and modulating inflammatory signaling. E2 and NO signaling decrease oxLDL internalization and oxidative stress through distinct endothelial pathways, while promoting endothelial activation and monocyte recruitment. At the same time, E2 attenuates macrophage inflammatory responses, indicating a dissociation between leukocyte recruitment and downstream inflammatory activation. Together, these results reveal both a context-dependent and context-independent role for E2 and NO signaling in regulating early atherosclerotic processes. Addressing these mechanisms will shed light on how estrogen-dependent pathways influence early disease progression, providing a mechanistic framework to inform future therapeutic strategies targeting vascular protection.

## Supporting information

Graphical Abstract

## FUNDING SOURCES

This work was supported by start-up funds provided to A.B.W. by High Point University.

## ACKNOWLEDGEMENTS

The authors thank the High Point University core facilities for technical support. We also acknowledge all members of the Wanschel laboratory for helpful discussions and assistance.

## CONFLICT OF INTEREST

The authors declare that the research was conducted in the absence of any commercial or financial relationships that could be construed as a potential conflict of interest.

## CRediT Authorship Contribution Statement

E.S., H.S., and G.A.M. performed experiments, analyzed data, and contributed equally to this work. C.P.S. and M.Z. contributed to data acquisition and analysis. M.H.K. contributed to in vivo studies and data interpretation. A.G.S. contributed to study design and critical data interpretation. A.B.W. conceived and supervised the study, secured funding, and wrote the manuscript. All authors critically revised the manuscript and approved the final version.

## DATA AVAILABILITY STATEMENT

The data needed to evaluate the conclusions of this study are present in the text, figures, and supplementary materials. Raw data may be obtained upon request.

## ABBREVIATIONS

CVD: Cardiovascular Disease
CAD: Coronary Artery Disease
Drp1: Dynamin-related Protein 1
E2: 17β-Estradiol
ED: Endothelial Dysfunction
eNOS: Endothelial Nitric Oxide Synthase
GDF15: Growth Differentiation Factor 15
HAECs: Human Aortic Endothelial Cells
HFD: High-Fat Diet
IL-1ra: Interleukin-1 Receptor Antagonist
LDLr^−/−^: Low-Density Lipoprotein Receptor Knockout
LOX-1: Lectin-like Oxidized LDL Receptor-1
MIF: Macrophage Migration Inhibitory Factor
MIP-3α: Macrophage Inflammatory Protein-3 Alpha
MMP9: Matrix Metalloproteinase 9
NO: Nitric Oxide
OPA1: Optic Atrophy 1
oxLDL: Oxidized Low-Density Lipoprotein
ROS: Reactive Oxygen Species
SNAC: S-nitroso-N-acetylcysteine
SR-BI: Scavenger Receptor Class B Type 1
SOD-2: Superoxide Dismutase 2

## Notes

### Competing Interest Statement

The authors have declared no competing interest.

### Summary of Updates

Expanded mechanistic interpretation of estrogen and nitric oxide signaling in endothelial lipid handling mitochondrial dynamics and inflammatory responses Added new analyses on sex specific differences in lesion formation blood pressure regulation and cardiac remodeling Included additional discussion on LOX1 caveolin1 and SRBI pathways Revised abstract introduction results and discussion for improved clarity and integration of in vitro and in vivo findings. Graphical Abstract added

