## Supplementary figures and images for "Estrogen-Nitric Oxide Signaling Modulates Mitochondrial Dynamics and Endothelial Lipid Handling to Protect Against Early Atherosclerosis"

### Graphical Abstract

**GRAPHICAL ABSTRACT**

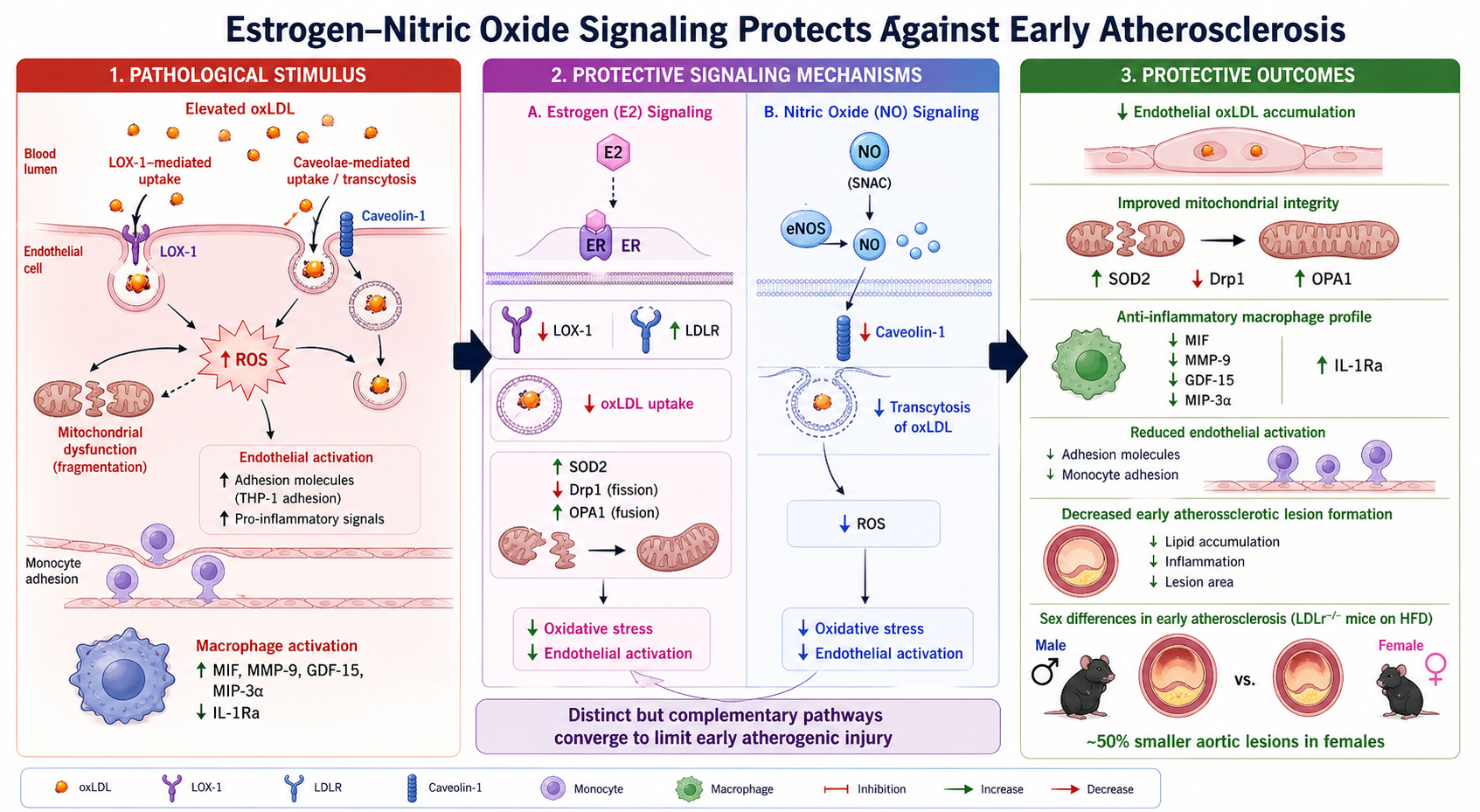
